# Patterned Metal Grids for Flexible and Transparent Neural Microelectrode Arrays

**DOI:** 10.1101/2023.05.08.539822

**Authors:** Ivânia Trêpo, Joana V. Pinto, Ana Santa, Maria E. Pereira, Tomás Calmeiro, Beatriz Coelho, Célia Henriques, Rodrigo Martins, Elvira Fortunato, Megan R. Carey, Hugo G. Marques, Pedro Barquinha, Joana P. Neto

## Abstract

Flexible and transparent microelectrodes can provide large-scale neural recordings with temporal and spatial resolution when used alongside functional calcium imaging. Patterned metal grids defined by direct laser writing (DWL) are a promising approach for these electrodes, as they resort to standard microfabrication processes and materials, allowing the possibility of mass production. For these reasons, a study exploring transparent grid-based electrodes using DWL for measuring electrocorticography signal was performed. Patterned metal grids with 1 μm of linewidth and 22 μm of spacing between lines showed a sheet resistance of 6 Ω/sq and a transmittance of 81% at 550 nm. The grids were transferred to a 5 μm Parylene-C membrane using an optimized procedure that involves an oxygen plasma pre-treatment. This procedure ensures mechanical robustness and stability of the grids. Finally, a flexible and transparent prototype was fabricated with a microelectrode array composed by 16 electrodes with 500 μm of diameter. These microelectrodes shown an impedance of 10 kΩ at 1 kHz in saline solution and they are highly conformal facilitating in vivo implantation and the recording of neural activity in the mouse cerebellum surface. To conclude, patterned metal grids based-electrodes exhibit a promising performance compared to transparent conductive oxides or graphene. Moreover, the introduction of DLW enables easy and fast manipulation of grid shape and dimensions without the need of physical masks, while keeping large scale compatibility, which is important for tools used in neuroscience community.

## 1. Introduction

Electrocorticography (ECoG) is an extracellular electrophysiological monitoring technique that uses microelectrode arrays (MEAs) placed on the brain surface to detect electrical signals generated from neural activity. Flexible ECoG MEAs can cover a large area of cortical surface, thus providing a view of distributed populations of neurons with sub-millisecond precision and minimal brain damage, but with limited spatial resolution [1,2]. Each electrode is sensitive to the activity of hundreds to thousands of neurons in its vicinity and to all transmembrane currents in the extracellular space (e.g., synaptic and action potentials currents)[3]. Alternatively, functional calcium imaging can be used to measure neural activity with single-cell resolution across large populations of neurons. With this technique neuroscientist can record the activity of genetically-targeted neurons using fluorescent calcium-sensitive indicators. [4]. However, these calcium signals are relatively slow and cannot provide precise information about the timing of neural activity relative to animal behavior [5].

In neuroscience, ECoG and functional calcium imaging are techniques normally used separately. Nevertheless, temporal and spatial resolution (i.e., to know when and where) are both relevant to understand interactions and dynamics of neural networks and to elucidate the neuronal mechanisms underlying brain function [6]. In this manner, transparent ECoG MEAs associated with functional calcium imaging can be used to obtain the necessary spatial and temporal resolution, covering the dynamics of many neurons, thus providing a detailed view of neural activity without penetrating the brain.

Firstly, to enable the combination of electrical and optical measurements, it is desirable that the ECoG substrate is transparent, conformable and flexible so that it can adapt itself to the curvilinear surface of the brain. Parylene-C is a common choice for such applications [7–12], since it is a FDA-approved biocompatible polymer [13]. In addition, the electrodes also need to be transparent in order to be used alongside functional calcium imaging, but they also need to be as conductive as possible. In such manner, there is an interplay between the electrical resistivity of the chosen electrode material and its optical transmittance. Nowadays, new materials, not originally developed for neural interfaces but for optoelectronics applications, are becoming promising candidates to sense neural signals. There have been various attempts to develop transparent ECoG MEAs using indium-tin oxide (ITO) [14,15], graphene [16,17] and conductive polymers [18–20]. ITO is known to be unsuitable for stable flexible and/or biological applications, due to its ceramic brittle nature. Besides, indium scarcity and pulmonary toxicity are additional concerns [21]. Graphene films are difficult to fabricate through reproducible and large-scale methods, hindering the possibility of mass production. Conductive polymers suffer from precarious electrical stability over time in biological mediums [22]. Furthermore, these materials need an extra metal layer for interconnections, which will add one more step in the fabrication process. Finally, they were not widely adopted by companies (and thus not used by neuroscientists) because the standardized protocols in place for neural electrodes are developed around inert and biocompatible metals, namely platinum, gold, and iridium.

Therefore, emergent transparent and flexible metallic nanostructures, such as metal nanowires, thin metal films and patterned metal grids (PMGs) are currently considered the best candidates due to their inherently high electrical conductivity, optical transparency, mechanical robustness, and cost competition [23]. Moreover, PMGs are advantageous for fabricating ECoG electrodes as they resort to standard microfabrication processes and materials, which allow for the production of large quantities at low-cost [23,24]. By using state-of-the-art techniques to pattern metals, namely direct laser writing (DLW), it is possible to explore optimal grid (i.e., with best compromise between transmittance and electrical resistance) and MEAs design in a fast and controllable manner, without imposing the fabrication of expensive physical masks.

This work reports on the study and optimization of PMG design for ECoG electrodes, first through simulation (COMSOL Multiphysics and Ansys Lumerical) and then through fabrication based on DLW. Throughout this work, gold PMG-based electrodes kept excellent optoelectrical properties, exhibiting a high optical transparency of 81% at 550 nm, and a sheet resistance of 6 Ω/sq. Furthermore, an ultra-thin and transparent ECoG prototype with 16 PMG-based electrodes was obtained through reproducible and controlled methods, demonstrating not only mechanical robustness through the stability of sheet resistance over the bending tests, but also low-impedance electrodes, which allowed to record neural electrical activity from the mouse’s surface brain.

## 2. Materials and Methods

### 2.1. PMG Design and Simulation

In order to assess the effect of PMG dimensions on their electrical and optical properties, simulations were made using COMSOL Multiphysics® software (AC/DC module) and Ansys Lumerical – FDTD (Finite Difference Time Domain) Solver, respectively. COMSOL simulations were based on a previous work [25]. An electrical potential was applied across the simulated PMG and the current density, J (A/m^2^), was calculated. Lastly, sheet resistance values were taken from simulations. A gold thickness of 80 nm was kept during all simulations, as well as a linewidth of 1 μm, only varying the spacing between lines (from 6 µm to 24 µm). Optical FDTD simulations were carried out using Lumerical software to assess the transmittance of each PMG. Since PMG structure is periodic, the FDTD region was reduced to only one period (one square) to reduce simulation time. For more details, consult Section 1 (PMG Sheet Resistance and Transmittance Simulations) of Supplementary Material.

### 2.2. PMG Fabrication

After proper glass substrate cleaning using acetone and isopropyl alcohol, each substrate was spin-coated with positive photoresist AZ® ECI 3012, from MicroChemicals GmbH, and then soft-baked. A tabletop μPattern Generator 101 Heidelberg Instruments Direct Laser Writing equipment was used to pattern the different grid designs (i.e., with different linewidth and spacing) onto the positive photoresist. This step was followed by a post exposure bake, and then immersing the sample in developer solution (AZ® 726 MIF – MicroChemicals GmbH), finishing with a post-development bake. Afterwards, a Ti/Au (6 nm and 80 nm, respectively) deposition was carried out by electron-beam evaporation at room temperature. The metal was later patterned by lift-off in acetone.

### 2.3. PMG Characterization

PMGs electrical characterization was performed in Nanometrics HL5500 Hall Effect Measurement Systems, through the Van der Pauw Method, in order to obtain sheet resistance values. Optical characterization took place in Spectrometer UV-Vis-NIR – Perkin Elmer Lambda 950 for transmittance measurements, using an integrating sphere, in the wavelength range of 250 to 1500 nm, with a step of 5 nm, and a gain of 2. Additional morphological characterization was performed through Scanning Electron Microscopy (SEM, Zeiss Auriga Crossbeam Microscope) and Atomic Force Microscopy (AFM, Asylum Research MFP-3D Stand Alone AFM System) in intermittent contact mode.

### 2.4. ECoG MEA Prototype Fabrication

A schematic illustrating the fabrication process is depicted in Figure 1.A. The substrate carriers for the prototype were 3-inch p-type silicon wafers. The deposition of Polyvinyl Alcohol (PVA) solution as a sacrificial water-soluble layer (Figure 1.A.1) and the deposition of Parylene-C, both as substrate (Figure 1.A.2) and as encapsulation (Figure 1.A.7) layers, were performed as reported by Neto et al. [7]. Parylene-C substrate and encapsulation layers have 5 µm and 1 µm of thickness, respectively. Photoresist patterning occurred as with the procedure previously described for the PMGs (Figure 1.A.3 and Figure 1.A.4). Afterwards, a 5-minute oxygen plasma pre-treatment was applied to the sample using Minilock – Phantom Reactive Ion Etching (RIE) from Trion Technology, to improve metal adhesion to Parylene-C. The next step was the Ti/Au deposition (Figure 1.A.5) followed by ultrasonic lift-off (Figure 1.A.6). Subsequently, in order to etch the encapsulation layer from the 16 electrodes and contact pads, a negative photoresist (AZ® nlof 2020 from MicroChemicals GmbH) was spin-coated and soft-baked (Figure 1.A.8). This photolithography process was made using a Karl-Suss MA6 UV Mask Aligner (Figure 1.A.9). After the post-exposure bake, Parylene-C etching took place in the RIE equipment using oxygen plasma (Figure 1.A.10). The process ended by peeling off the samples from the substrate (Figure 1.A.11).

**Figure 1.**
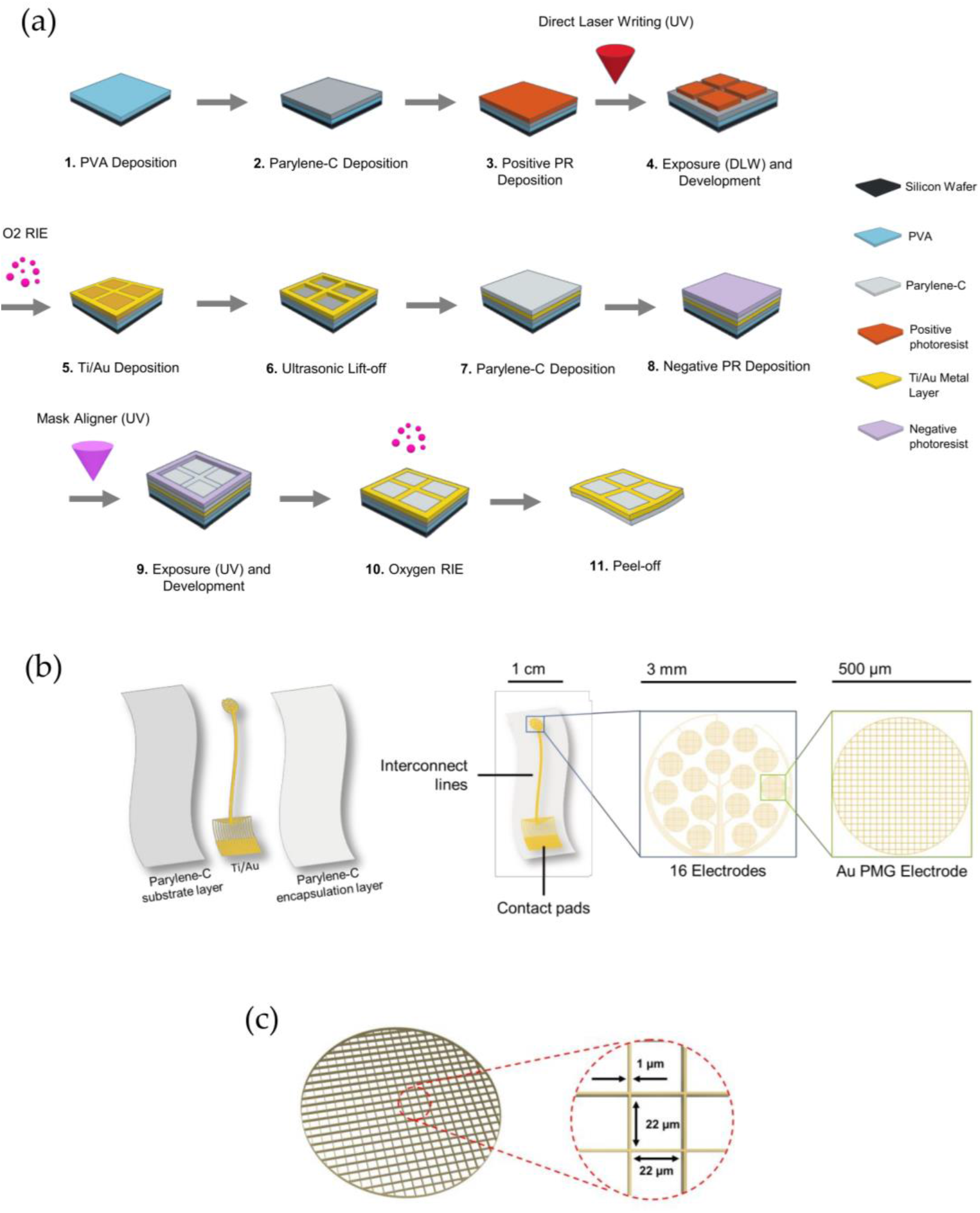
ECoG MEA with gold PMG fabrication schematic (illustrative and not to scale). A) Step-by-step fabrication of the prototype is depicted. B) Schematic overview of the device where different layers, different parts and their dimensions are described. C) Individual Au PMG-based electrode with 1 µm of linewidth and 22 µm of spacing indicated.

In Figure 1.B, the 3 main layers of the prototype are shown, followed by an illustration of the dimensions of each part of the produced ECoG device. The entire prototype has around 3 cm of length and consists of three main parts: the ‘head’ of the device with 16 electrodes, the interconnect lines, and the output pads to connect to a Zero Insertion Force (ZIF) adapter. The ‘head’ of the device has a circular shape with 3 mm of diameter because of the limitation imposed by the cranial window size used in calcium imaging setup, which is equally 3 mm wide, thus taking advantage of the available area. The MEA has 16 electrodes with a diameter of 500 μm distributed over 3 mm. The final electrodes were fabricated considering the gold PMG with 22 μm of spacing and 1 μm of linewidth (Fig 1.C.).

### 2.5. ECoG MEA Prototype Characterization

Prototypes were then characterized by connecting them to an adapter equipped with a ZIF and Omnetics connector, compatible with the measuring equipment, which comprises the Open Ephys acquisition board along with the RHD2000 series interface chip that amplifies and digitally multiplexes the signal from the 16 extracellular electrodes (Intan Technologies). The impedance magnitude at 1 kHz of each electrode was measured using a protocol implemented by the RHD2000 series chip from Intan, with a two-electrode cell configuration: the electrodes of the ECoG device and the reference electrode, an Ag/AgCl wire (Science Products GmbH, E-255), were immersed in saline solution (0.9% NaCl solution). Moreover, we have performed a preliminary experiment in vivo to evaluate the ECoG surgery implantation protocol and its compatibility with the surgical procedures used for calcium imaging. A male C57BL/6 mouse was anesthetized with isoflurane and head-fixed using a stereotaxic frame. We performed a craniotomy around the center of the occipital using a 3-mm biopsy punch aiming towards the cerebellum. The transparent ECoG MEA was placed in the center of the craniotomy and covered with a circular glass transparent coverslip (3 mm diameter), which was then pushed into the craniotomy and glued into the skull. Then, a customized steel headplate was secured on the skull of each animal with dental cement (Super-Bond C&B). The animal was head-fixed on a rotary self-paced treadmill with 250 mm diameter. Recordings were conducted using and OpenEphys board with 32 channels, in a frequency band of 0.1 – 7,500 Hz, sampled at 30 kHz with 16-bit resolution and were saved in a raw binary format for subsequent offline analysis. All procedures were carried out following the European Union Directive 86/609/EEC and approved by the Champalimaud Centre for the Unknown Ethics Committee and the Portuguese Direção Geral de Veterinária (Ref. No. 0421/000/000/2020).

## 3. Results and Discussion

### 3.1. PMG Geometry

Several PMG geometries have been reported in literature, not originally for neural interfaces but mainly for optoelectronic applications. An overview of the most used metal grid designs, their fabrication method, and their properties can be consulted in Section 2 of Supplementary Material. The main objective has been to find the best compromise between optical transmittance and sheet resistance, while assuring good process yield. The most studied patterns are square grids [26–30], honeycomb structures [31–33], and circular shapes [25,34–36]. In Figure 2, for the same area and considering a linewidth of 1 µm and a spacing of 10 µm between lines, circular shapes have 38% of area coverage (Figure 2.A), while honeycomb structures have 20% (Figure 2.B), and square grids have the lowest area coverage of 18% thus increasing optical transparency (Figure 2.C). Our work took into consideration this analysis and it was carried out considering the square grid geometry to maximize optical transmittance.

**Figure 2.**
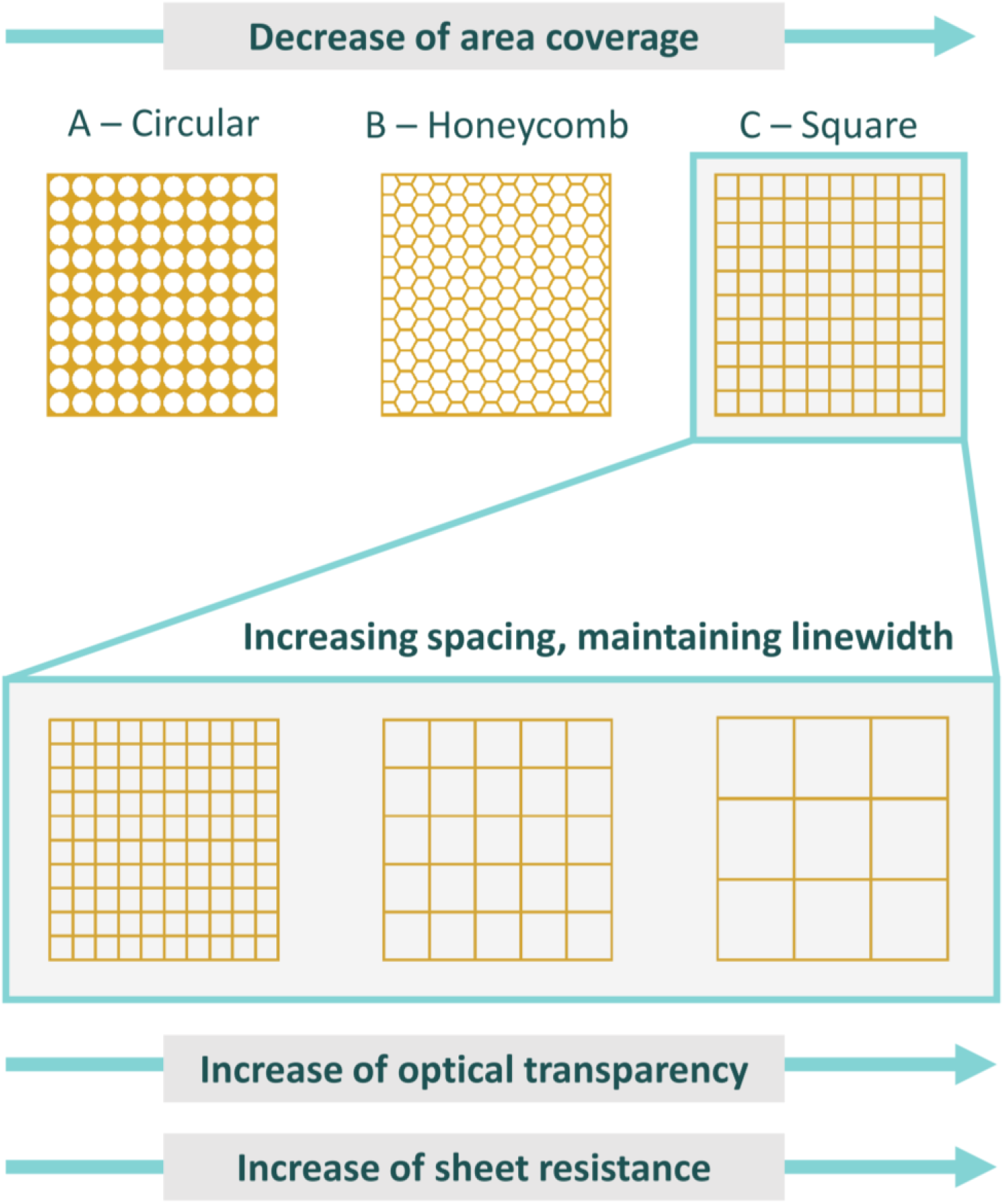
Influence of metal grid geometry in area coverage, considering a linewidth of 1 µm and a spacing of 10 µm between lines: A) Circular shape, B) Honeycomb structure and C) Square grid. Schematic of metal grid geometries and dimensions, and their influence on optoelectronic properties, as well as the trade-off between spacing and linewidth to obtain optimal optoelectronic properties.

### 3.2. PMG Electrical and Optical Characterization

Figure 3.A depicts the relation between optical transmittance and sheet resistance of the PMGs, on glass, for a fixed linewidth of 1 µm and for different line spacing (between 6 µm and 24 µm). A good agreement between simulation and experimental data is observed, concerning the trends of optical transmittance and sheet resistance with increasing PMG spacing (for more information see Section 3 of Supplementary Material). Nevertheless, the absolute values of the two data series (simulation and experimental) exhibit some offset. Regarding sheet resistance, the average difference between theoretical and experimental results is 14 ± 6 %, whereas regarding transmittance this difference is 12 ± 3 %. The deviation between simulation and experimental results may be due to: i) a difference in PMG dimensions between simulation and fabricated grids for characterization, ii) defects in PMG, which emerged during fabrication, and iii) the use of simplified models for simulation on behalf of computational time, which may not reflect with full fidelity the behaviour of PMG. Despite the discrepancies between experimental and simulation data (less than 20%), both follow a well-matched trend. As shown in Fig 3.A. the density of the fabricated PMGs diminishes as the spacing between lines increases, thus resulting in a rise in transmittance values. However, fewer lines lead to fewer pathways for electron flow, which results in an increase of the sheet resistance, as expected. When selecting the proper PMG spacing there is also another parameter to consider: the number of lines within the electrode, which in this work is 500 μm of diameter. If there is not a sufficient number of lines in the electrode, then the electrode impedance will be high. The selected value of spacing must also stand within the range of minimum transmittance of 80% (at 550 nm) to ensure the electrode has high optical transparency. The first PMG design fitting this criterion was the one with 22 μm of spacing, showing a sheet resistance of 6 Ω/sq. Therefore, 22 μm of spacing was the optimal compromise found among these three criteria, which are: low sheet resistance, high optical transparency, and a low impedance value indirectly implied by the number of lines within the electrode. The optoelectrical values of the resulting PMG (1 µm of linewidth and 22 µm of spacing) placed our work in a prominent position regarding previous studies with other transparent electrode materials for electrophysiological recordings, as shown in Figure 3.B. This is a direct result of tuning the dimensions of the PMG through DLW, so that optimal values are attained, with the potential of further optimization, reaching a successful compromise between optical and electrical properties as aforementioned.

**Figure 3.**
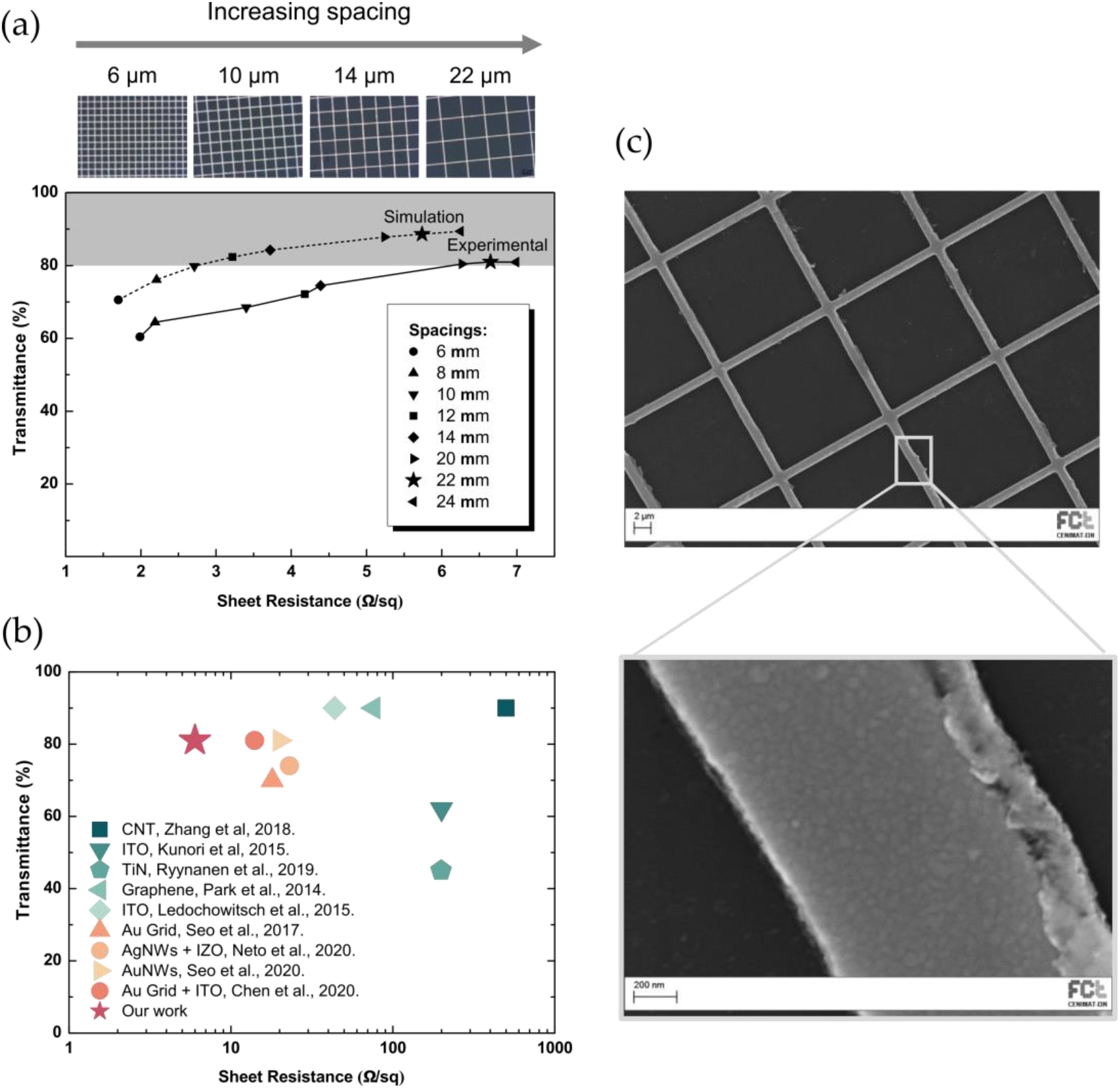
PMG characterization. A) Graphical comparison between simulation and experimental data for a fixed linewidth of 1 µm, varying the spacing between grid lines. B) Positioning of the obtained optoelectrical experimental results on PMGs with respect to literature focusing on transparent ECoG electrodes. C) SEM micrograph of Au PMG with a close-up of PMG partial line where surface morphology is evident.

The PMG, and a close-up view of a PMG partial line can be seen in Figure 3.C. Despite some resist residuals that reveal the lift-off process still requires additional improvements, very good pattern definition is obtained even using a very small linewidth (1 µm) with DLW. Moreover, in order to assess how accurately PMG features were reproduced during photolithographic processes, the proximity effect was analysed, and it showed to be negligible for the linewidth and range of the line spacing considered (see Section 4 of Supplementary Material). It is noteworthy that different gold thicknesses were considered during COMSOL simulations (Section 5 of Supplementary Material) and, similarly as reported by Obaid et al. [37], sheet resistance did not vary significantly considering the range of 60 to 120 nm of Au thickness. Therefore, a gold thickness of 80 nm was selected to facilitate the lift-off step and reduce production costs, without compromising the sheet resistance value at the end.

### 3.3. Gold PMGs on Parylene-C

This section is dedicated to the production of flexible gold PMG on an ultrathin Parylene-C membrane. The yield on untreated Parylene-C substrates was found to be very low. Therefore, in order to improve the fabrication yield on Parylene-C substrate, a treatment with oxygen plasma was evaluated before the metal deposition. AFM analyses were conducted and the variation of surface roughness over oxygen plasma time is evident, as seen in the results presented in Figure 4.A. Depending on the time oxygen plasma is applied, it starts to gently etch Parylene-C, increasing surface roughness [38]. The Ti-Au layer, deposited onto the Parylene-C membrane, will then acquire its surface roughness. The RMS values (in nm) are obtained from the analysis of the surface height variations. With no oxygen plasma pre-treatment (i.e., 0 min.) the RMS value is 8.4 nm, whereas after 5 minutes of oxygen plasma the RMS value is 71.7 nm. The 5 minutes treatment showed to be sufficient to enhance the mechanical stability of the prototype. Moreover, for this specific application, surface roughness is welcomed since it decreases electrode impedance. Nevertheless, the plasma treatment also resulted in the undesired removal of some photoresist from the sidewalls of the patterned structures. This is due to the isotropic component of the oxygen dry etching. In such manner, linewidth can slightly increase due to this intermediate step. It was concluded that linewidth increases approximately 27% after oxygen plasma pre-treatment in patterned photoresist. A microscope image illustrating the effect of oxygen plasma to the linewidth can be found in Section 6 of Supplementary Material.

**Figure 4.**
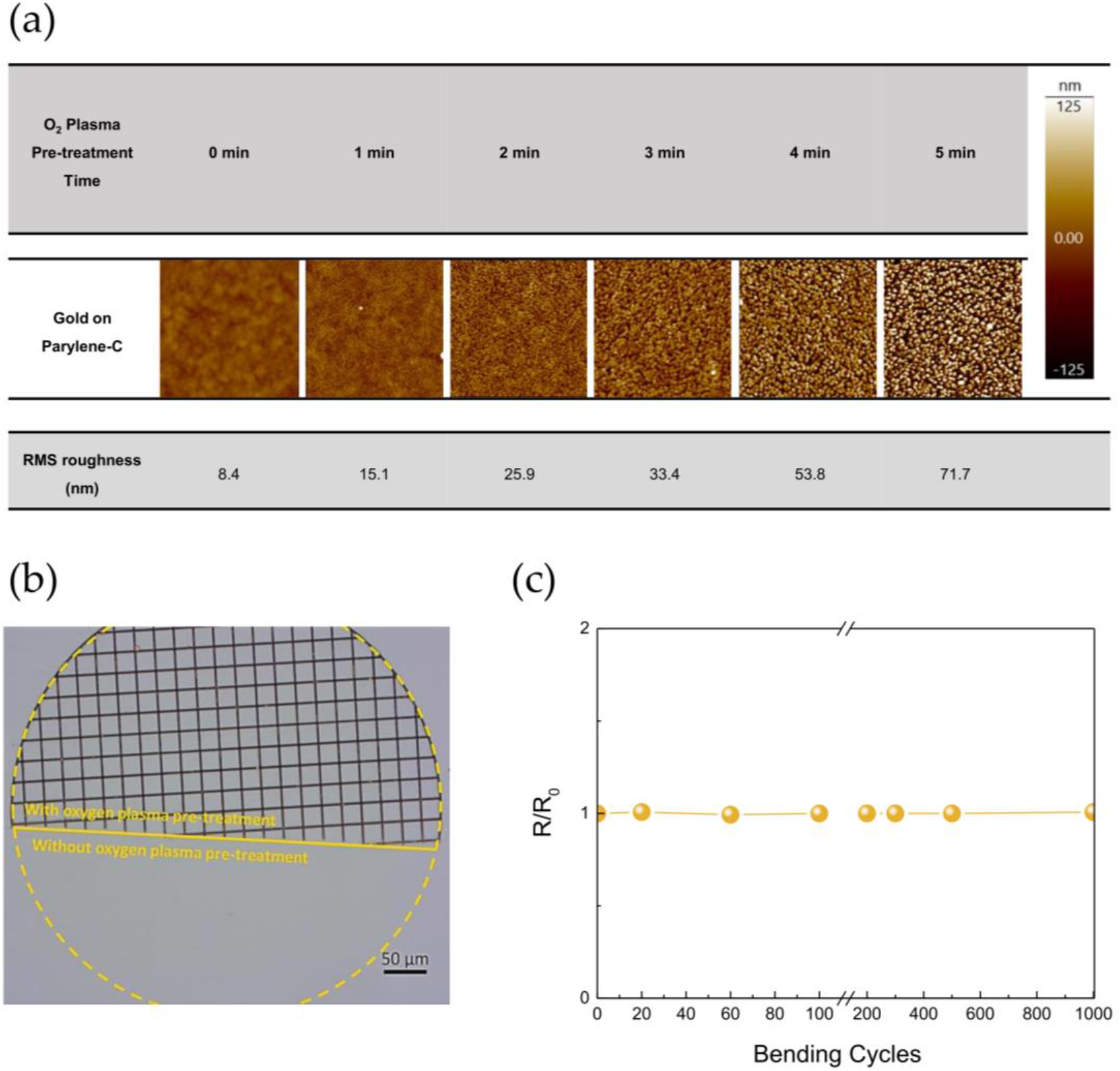
Effect of oxygen plasma pre-treatment on Parylene-C membrane for Au PMG fabrication. A) AFM scans showing oxygen plasma pre-treatment effects on gold on Parylene-C surface roughness. Scan size is 5 µm × 5 µm and the color bar indicates the surface height. B) Microscope image showing that without this step (5 min O_2_ plasma) the gold PMG does not adhere to the substrate after lift-off. C) Testing sheet resistance variation of gold PMG during bending cycles. The ratio R/R0 expresses the sheet resistance at a certain bending cycle (R) with respect to the initial sheet resistance (R0).

It is possible to observe in Figure 4.B that the sample region that was not exposed to oxygen plasma pre-treatment is completely absent after the lift-off due to the lack of adhesion, while the region submitted to this treatment is uniform, which indicates that this step is crucial during PMG MEA manufacturing process. Thus, this oxygen plasma step promotes polymer-metal adhesion [39,40], improving the prototype robustness and the process yield.

In order to determine the mechanical stability of the flexible PMG, the sheet resistance was evaluated after performing repeated bending cycles with ∼2.5 mm of bending radius. The change ratio (R/R_0_) in PMG was recorded as a function of the bending cycle and it is depicted in Figure 4.C. If this ratio is superior to one, then the R (sheet resistance at a certain bending cycle) is larger than R_0_ (initial sheet resistance), indicating a decrease in the electrical stability of the PMG. Nonetheless, the results demonstrate that the sheet resistance of the PMG with 22 µm of spacing have negligible changes after consecutive bending cycles, thereby exhibiting a stable electrical performance even after more than 1000 bending cycles. This is due not only to the ductility of gold, but also to the previous oxygen plasma pre-treatment which provided mechanical robustness to the prototype.

### 3.4. Cytotoxic Effects Assessment of Parylene-C

The materials used in the device, namely Parylene-C, gold and titanium, are inert and biocompatible. Nevertheless, to evaluate if the Parylene-C after plasma pre-treatment present toxicity and interact negatively once implanted in the brain, cytotoxic effects were analyzed in vitro according to the International Standard (ISO 10993–5) using the extract method. The complete protocol can be consulted in Supplementary Material – Section 7. The results show that the relative viability obtained for cells exposed to the extracts from the untreated and from the oxygen plasma pre-treated Parylene-C membranes are 99.5 ± 2.6 % and 100.7 ± 3.5 %, respectively. It is possible to infer through these results that the developed PMGs on Parylene-C do not bring any cytotoxic effects, regardless of whether Parylene is treated or not.

### 3.5. ECoG MEA Prototype Characterization

After PMG fabrication and characterization in Parylene-C flexible substrate, prototype production followed as described in the “2. Materials and Methods” Section. The ultrathin Parylene-C membrane as substrate enables a highly-conformable prototype as shown in Figure 5.A. The prototype conforms to the surface of a delicate and soft petal. Impedance analysis at 1 kHz is a common metric to determine how electrodes will perform in vivo. In Figure 5.B, it is possible to observe the impedance magnitude measured at 1 kHz in saline solution for 7 prototypes and their individual electrodes. The mean ± standard deviation value for impedance, for all the prototypes fabricated, was 10.2 ± 3.2 kΩ, and for the control prototype (whose electrodes consist of a plain thin film of gold with 80 nm of thickness, instead of a gold PMG) was 2.1 ± 0.1 kΩ. In this manner, authors conclude that low-impedance transparent PMG-based electrodes on flexible and ultrathin substrate were achieved, using reproducible procedures. The Figure 5.C shows the microscope image of one ECoG MEA prototype, where some of the transparent PMG-based electrodes are visible, with one of them being denoted with a white dashed line. The image on the right shows a magnified view of a single gold PMG-based electrode with 500 µm of diameter.

**Figure 5.**
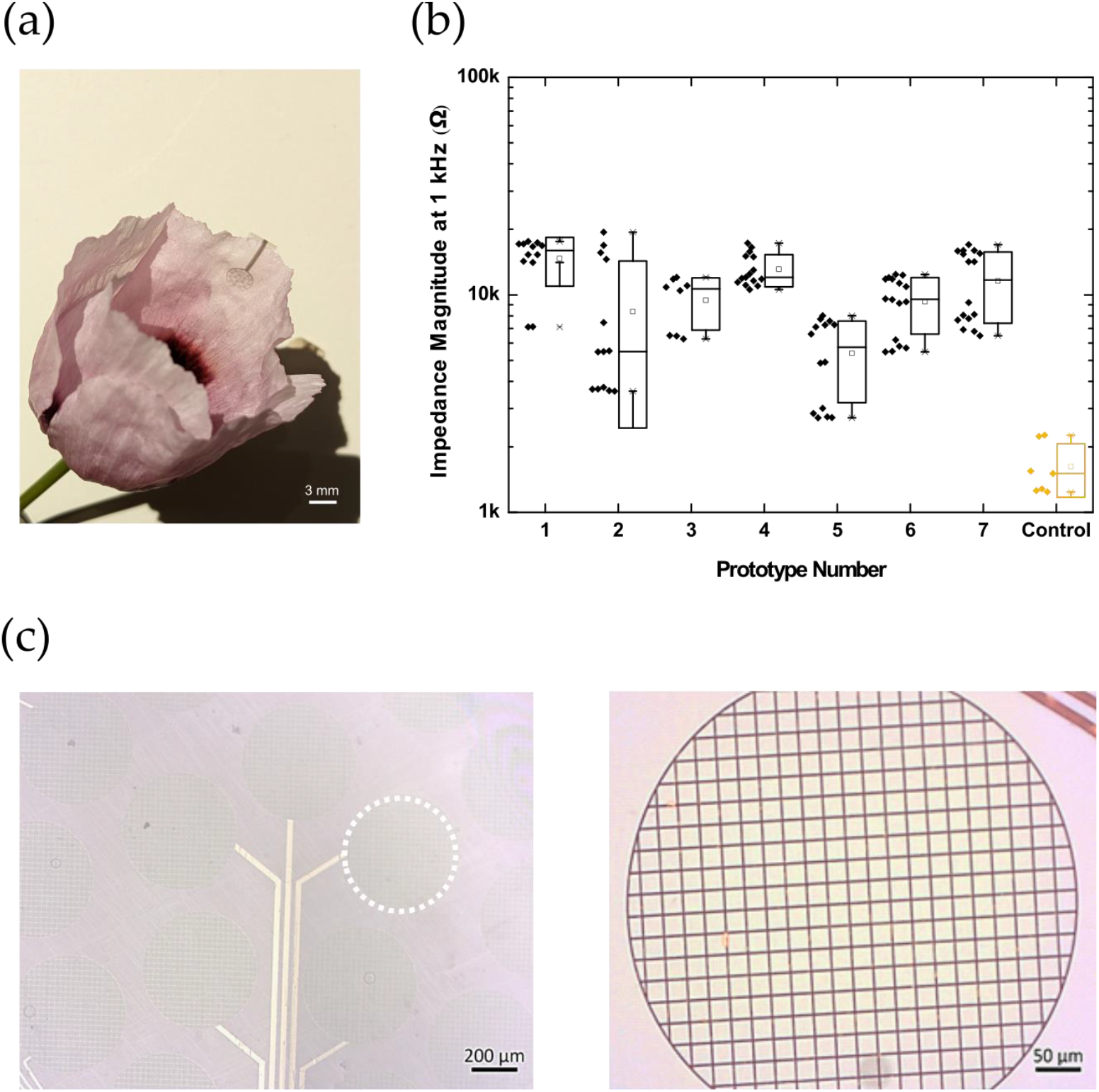
Transparent and flexible ECoG prototype characterization. A) The ultrathin transparent ECoG conforms to the surface of a poppy petal. B) Impedance magnitude measured at 1 kHz in saline solution as a function of the various electrodes and prototypes. The control prototype electrodes consist of a thin film of gold with 80 nm of thickness. In the box plots: the line is the median, the square is the mean, the box 1st to 3rd quartile, and whiskers is the 1.5 × interquartile range above and below the box. C) Microscope image of the 16 electrodes with their respective interconnect lines, and an inset of one Au PMG-based electrode.

Figure 6.A illustrates the device positioned in a mouse brain, with a close-up image of the electrode array top view. A preliminary acute experiment was used to evaluate the ECoG surgery implantation protocol and its compatibility with the surgical procedures used for calcium imaging. Thus, an ECoG prototype was placed onto the surface of cerebellum through the cranial window, and after it was covered and fixed under a glass window. The electrode array and craniotomy limits are indicated with a black dashed line. It is possible to observe the brain surface underneath, as well as cerebral vasculature through the prototype suggesting that this electrode array may be of great use to perform functional calcium imaging. Additionally, the low thickness of the device (∼6 µm thick) was a key factor for a stable and intimate electrode-tissue coupling between prototype and brain. Additionally, very preliminary ECoG data was collected from cerebellum while the animal was walking freely on a treadmill. As an example, Figure 6.B shows the electrical signals recorded in three electrodes of the array. The correspondent power spectra plot (Figure 6.C) of ECoG signals extended up to high-frequencies. It was previously reported that ECoG microelectrodes are able to record cerebellar cortical activities in the rat, both local field potential (LFP) and multiunit activity (MUA) [41]. This suggests that our ECoG PMG-based electrodes are able to record high quality signals, sampling slow synaptic activity as LFP and fast action potentials as MUA, due to the distance and large area electrodes [42,43].

**Figure 6.**
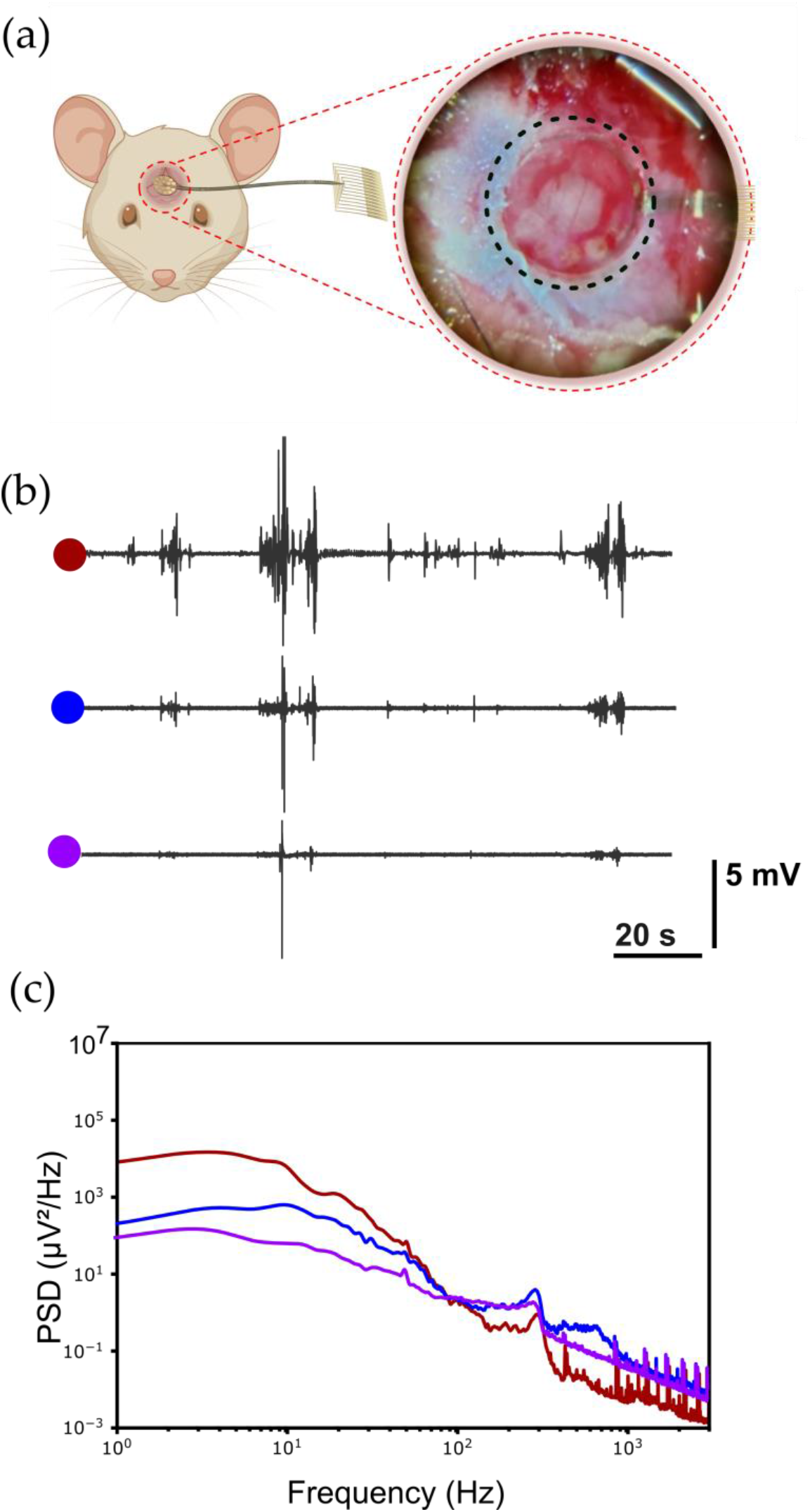
ECoG recordings from cerebellum while the mouse was on a treadmill. A) Scheme of ECoG prototype with 16 gold PMG-based electrodes implanted on the mouse’s brain surface. Inset: ECoG laying on the cerebellum of a mouse through a cranial window. B) Traces (2 min long) of recorded neural activity at cerebellum surface from three electrodes of the MEA. The red, blue and pink circle identify the color of each electrode. C) Signal power spectra density for the three ECoG signals (raw data). For the calculation of the power spectral density we used Welch’s method from SciPy package available at https://docs.scipy.org/doc/scipy/reference/generated/scipy.signal.welch.html, selecting nfft as data length, nperseg as sampling frequency*0.5 and noverlap as nperseg *0.5. The sampling frequency was 30,000 Hz.

Finally, Table 1 outlines the transmittance, sheet resistance and impedance values, electrode materials and respective dimensions to illustrate the interplay between those parameters of PMG-based ECoG electrodes. We benchmarked the performance of the fabricated PMG electrodes in comparison to other developed transparent ECoG electrodes. The obtained results from this study placed us in a prominent position regarding the current state-of-the-art. Moreover, the bottleneck for several approaches is the fabrication efficiency, reliability, material cost and compatibility with large-scale processes. PMGs fabricated with DLW are advantageous for fabricating ECoG electrodes because (i) PMGs use standard fabrication processes and materials, which allows for the fabrication of large quantities at low cost, (ii) DWL enables changing grid design in a fast and controllable manner when neuroscientists need to, (iii) gold is biocompatible, highly conductive and ductile enabling stable and flexible electrodes, (iiii) when high density MEAs are envisaged, the high electrical conductivity of metals enables more efficient miniaturization when compared to other potential ECoG materials, such as transparent conductive oxides, and (v) the materials used for PMGs can also be used for MEAS interconnections, thus avoiding interfaces between dissimilar materials. Our work demonstrates an ultrathin and transparent metal-only based ECoG MEA that can be readily up-scaled and easily adapted to application-driven requirements of optical and electrical properties by adapting the design of the PMG

**Table 1.** ECoG transparent electrodes, their dimensions, impedance magnitude (at 1 kHz in saline solution), sheet resistance and transmittance values at 550 nm from previous works compared to the present study. (NA: Not Available; ⦰: Diameter).

| Reference | Electrode Material | Electrode Dimension | $\vert Z\vert $ @ 1 kHz | Sheet Resistance ($\Omega/\text{sq}$) | Transmittance @ 550 nm |
| --- | --- | --- | --- | --- | --- |
| [16] | Doped Graphene | 50 x 50 $\mu\text{m}$ | ~ 500 kΩ | NA | 90 % |
| [44] | Graphene | 100 x 100 $\mu\text{m}$ | 15 kΩ | NA | 50 % |
| [45] | Graphene | 200 $\mu\text{m}$ ∅ | 243 kΩ | 76 | 90 % |
| [17] | Graphene | 100 x 100 $\mu\text{m}$ | > 963 kΩ | NA | 90 % |
| [46] | CNT | NA | 0.2 MΩ | 150 – 500 | 70 – 90 % |
| [20] | PEDOT:PSS | 30 $\mu\text{m}$ | 46.5 kΩ | NA | 93 % |
| [47] | TiN | 30 $\mu\text{m}$ ∅ | 510 – 590 kΩ | 41 – 199 | 18 – 45 % |
| [14] | ITO | 250 $\mu\text{m}$ ∅ | 10 – 50 kΩ | 200 | 62 % |
| [48] | ITO | 80 $\mu\text{m}$ ∅ | 250 kΩ | NA | 80 % |
| [15] | ITO | 500 $\mu\text{m}$ ∅ | 300 kΩ | 43.7 | 90 % |
| [7] | IZO 200 nm | 500 $\mu\text{m}$ ∅ | 40.7 kΩ | 25 | 76 % |
| | AgNWs + IZO | 500 $\mu\text{m}$ ∅ | 23 kΩ | 23 | 74 % |
| [35] | Au Grid + PEDOT | 80 x 80 $\mu\text{m}$ | 11 kΩ | NA | 70 % |
| [12] | Au Grid | 80 x 80 $\mu\text{m}$ | 127 kΩ | 18 | 70 % |
| [49] | AuNW | 200 $\mu\text{m}$ ∅ | 33.9 kΩ | 3 - 20 | 81 % |
| [30] | ITO + Au PMG | NA | 5.4 – 18.4 kΩ | 5.6 – 14.1 | 59 – 81 % |
| **Our work** | **Au PMG** | **500 $\mu\text{m}$ ∅** | **10 kΩ** | **6** | **81 %** |

## 4. Conclusions

The field of neuroengineering has been revolutionizing neuroscience research and clinical practice. The development of transparent ECoG MEAs may be central for advancing neuroscience research by enabling recordings of a large neural population with high spatial and temporal resolution, increasing our understanding of the nature of the recorded signal.

In this project, we developed a flexible and ultrathin ECoG prototype with transparent electrodes. The key elements of these electrodes were (i) the gold patterned metal grid whose dimensions were freely adjusted through direct laser writing in order to obtain optimal optoelectrical properties and (ii) the 5 µm thick Parylene-C membrane as substrate. The optimized Au PMG, with 1 μm of linewidth and 22 μm of spacing, experimentally demonstrated very high optical transmittance (81% at 550 nm) and low sheet resistance (6 Ω/sq) values, thus matching the simulations made in firsthand. The prototype is composed by a microelectrode array with 16 electrodes (500 μm of diameter each distributed over 3 mm) with an impedance value of 10 kΩ at 1 kHz in saline solution. The prototype (∼6 µm thick) is mechanically robust due to an oxygen plasma pre-treatment of Parylene-C substrate. Moreover, when comparing the fabricated device with current literature, it presents very promising results, placing this work in a prominent position, enabling a new generation of transparent ECoG MEA based on state-of-the art techniques to pattern metal, which are reliable and compatible with large-scale production. In the future, this concept can be extended into sub-µm linewidths and spacing while keeping flexible substrate and large area compatibility, by using techniques such as Nanoimprint Lithography. This will allow to reduce electrode size and thus increase the number of electrodes within the device to attain high-density recording, as well as gaining the ability to record individual neuron’s action potentials, since electrodes are scaled down to cellular dimensions (i.e., ≤50μm)[50].

## Supplementary Materials

The following supporting information can be downloaded at: www.mdpi.com/xxx/s1, Section 1 – PMG Sheet Resistance and Transmittance Simulations; Section 2 – PMG Geometry Literature Overview; Section 3 – Simulation vs. Experimental Results; Section 4 – Optical Proximity Effect on PMG Lines; Section 5 – PMG Sheet resistance vs. Gold Thickness – COMSOL Simulations; Section 6 – PMG Linewidth after Oxygen Plasma Pre-treatment; Section 7 – Cytotoxic Effect Study Protocol (Materials and Methods).

## Author Contributions

Ivânia Trêpo: Conceptualization, Methodology, Investigation, Formal Analysis, Writing – Original Draft, Visualization. Joana V. Pinto: Methodology, Writing – Review & Editing, Visualization. Ana Santa: Methodology, Investigation. Maria E. Pereira: Methodology, Writing – Review & Editing. Tomás Calmeiro: Investigation. Beatriz Coelho: Writing – Review & Editing, Visualization. Célia Henriques: Resources, Investigation. Rodrigo Martins: Funding Acquisition, Writing – Review & Editing. Elvira Fortunato: Funding Acquisition, Writing – Review & Editing. Megan R. Carey: Resources, Writing – Review & Editing. Hugo G. Marques: Supervision, Conceptualization, Methodology, Investigation, Writing – Review & Editing. Pedro Barquinha: Supervision, Conceptualization, Methodology, Writing – Review & Editing, Project Administration, Funding Acquisition. Joana P. Neto: Conceptualization, Methodology, Investigation, Writing – Original Draft, Writing – Review & Editing, Visualization.

## Data Availability Statement

We encourage all authors of articles published in MDPI journals to share their research data. In this section, please provide details regarding where data supporting reported results can be found, including links to publicly archived datasets analyzed or generated during the study. Where no new data were created, or where data is unavailable due to privacy or ethical restrictions, a statement is still required. Suggested Data Availability Statements are available in section “MDPI Research Data Policies” at https://www.mdpi.com/ethics.

## Abbreviations

AFM: Atomic Force Microscopy
DLW: Direct Laser Writing
ECoG: Electrocorticography
EIS: Electrochemical Impedance Spectroscopy
FDTD: Finite Difference Time Domain
ITO: Indium Tin Oxide
MEA: Microelectrode Array
PMG: Patterned Metal Grid
PR: Photoresist
PVA: Polyvinyl Alcohol
RIE: Reactive Ion Etching
RMS: Root Mean Square
SEM: Scanning Electron Microscopy
ZIF: Zero Insertion Force

## Disclaimer/Publisher’s Note

The statements, opinions and data contained in all publications are solely those of the individual author(s) and contributor(s) and not of MDPI and/or the editor(s). MDPI and/or the editor(s) disclaim responsibility for any injury to people or property resulting from any ideas, methods, instructions or products referred to in the content.

