## Supplementary Materials for "Patterned Metal Grids for Flexible and Transparent Neural Microelectrode Arrays"

**List of Contents:**

**Section 1 –** PMG Sheet Resistance and Transmittance Simulations
**Section 2 –** PMG Geometry Literature Overview
**Section 3 –** Simulation vs. Experimental Results
**Section 4 –** Optical Proximity Effect on PMG Lines
**Section 5 –** PMG Sheet resistance vs. Gold Thickness – COMSOL Simulations
**Section 6 –** PMG Linewidth after Oxygen Plasma Pre-treatment
**Section 7 –** Cytotoxic Effect Study Protocol (Materials and Methods)

### S1 – PMG Sheet Resistance and Transmittance Simulations

In order to assess the dimensions that would offer an optimal trade-off between sheet-resistance and transmittance, we started by simulating PMG properties in COMSOL Multiphysics (sheet resistance) and in Ansys Lumerical – FDTD Solver (transmittance).

In **Figure S1.A**, taken from COMSOL software, we can see the current flow through the PMG. While horizontal lines are in bright orange, representing very high current density, vertical lines are dark blue, illustrating a low current flow. This is due to the fact that, in the simulation, the voltage was applied on the first vertical line of the PMG on the left, as shown in **Figure S1.A**. Regarding how optical FDTD simulations were made (**Figure S1.B**), since we have applied a periodic condition due to the periodic nature of the PMG, considering the period as one square, the FDTD region will only assume the area of that square, resulting in shorter simulation times.

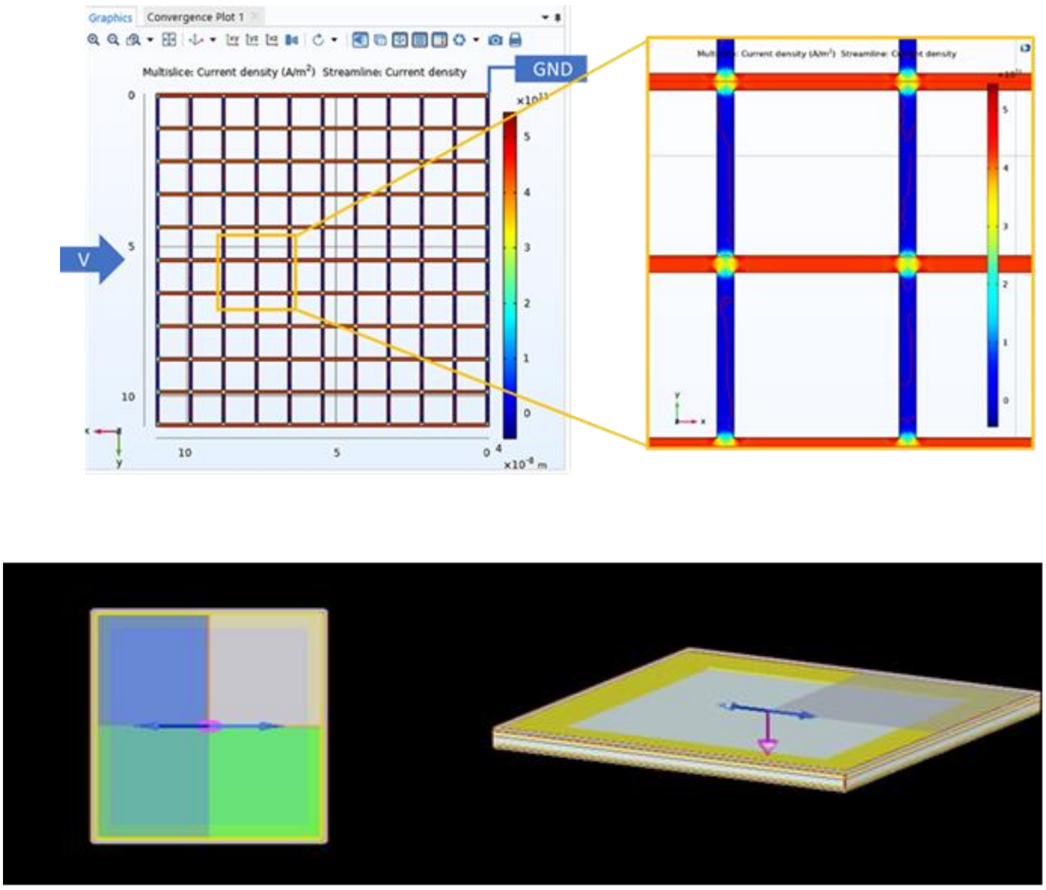


**Figure S1.** – PMG sheet resistance and transmittance simulations. Top:) Illustration of PMG simulations in COMSOL Multiphysics. Current density represented in false colors. Bottom: Schematic of the PMG simulation structure on Ansys Lumerical – FDTD Solver, with XY view (left) and perspective view (right).

**S2 – PMG Geometry Literature Overview**

A brief literature overview regarding the transmittance and the sheet resistance of the different PMG geometries was made and the results are contemplated in the **Table S2**.

**Table S2.** – Overview of the most used metal mesh geometry (square grids, honeycomb structures and circular shapes), their fabrication methods, and their properties (sheet resistance and transmittance considering material thickness). NA – Not Available.

| Geometry | Material | Technique | Thickness | Rs (Ω/sq) | T (%) | Reference |
|---|---|---|---|---|---|---|
| **Square** | Ag | Photolithography | 50 nm | 13.27 | 81.1 | [1] |
| | Ag | Selective metal condensation | 90 nm | 20 | 77 | [2] |
| | Au | NanoDrip | 200 nm – 1.5 µm | 8 | 94 | [3] |
| | Au/ITO | Photolithography | 80 nm | 14.1 | 81 | [4] |
| **Honeycomb** | Ag | Direct Laser Writing | 2.5 µm | 10 | 90 | [5] |
| | ITO/Ni/Ag/Al | Photolithography | 1 µm | <1 | NA | [6] |
| | Ag/Ni | Printing/Electroplating | 1.8 µm | 2.1 | 88.6 | [7] |
| **Circular** | Ag | Colloidal lithography | 17 nm | 12.8 | 72 | [8] |
| | Au | Nanoimprint Lithography | 40 nm | 2.12 | ~50 | [9] |
| | Au/IrO | Nanosphere lithography | 15 nm | NA | 70 | [10] |
| | Au/PEDOT | Nanosphere tolithography | 25 nm | NA | 70 | [11] |

#### S3 – Simulation *vs.* Experimental Results

In this section, it is possible to see all of the obtained results regarding transmittance and sheet resistance measurements, either by simulation or experimentation.

**Table S3.** – Optical and electrical PMGs characterization obtained through experimentation (Lab) and through simulation (Sim), considering an increase in spacing (from 6 μm to 24 μm, with a constant linewidth of 1 μm).

| Spacing (μm) | $R_S$ (Ω/sq) | | T(%) | |
|---|---|---|---|---|
| | Lab | Sim | Lab | Sim |
| **6** | 1.99 | 1.70 | 60.40 | 70.56 |
| **8** | 2.19 | 2.21 | 64.44 | 76.11 |
| **10** | 3.40 | 2.71 | 68.48 | 79.78 |
| **12** | 4.18 | 3.22 | 72.14 | 82.35 |
| **14** | 4.39 | 3.72 | 74.50 | 84.25 |
| **20** | 6.27 | 5.24 | 80.4 | 87.81 |
| **22** | **6.65** | **5.74** | **81.01** | **88.61** |
| **24** | 6.99 | 6.25 | 80.91 | 89.41 |

### S4 – Optical Proximity Effect on PMG Lines

In order to assess how accurately PMG features were reproduced during photolithographic processes, we studied the proximity effect on PMG lines. Considering the 10 µm PMG, since the lines are very close to each other, it was expected that the proximity effect was slightly more accentuated than the PMG with 22 µm of spacing. In this case, grid lines are more distant from each other, therefore a smaller proximity effect was foreseen. In the end, proximity effect for PMG lines with such dimensions was negligible. For this study, linewidth measurements were made using optical microscopy. The results are shown in **Table S4**.

**Table S4.** – Optical proximity effect analysis in positive photoresist (PR) patterned grid and gold PMG (20 measurements per image) for 1 µm of linewidth. The images were obtained through optical microscopy.

| Spacing | PR PMG | | Au PMG | |
|---|---|---|---|---|
| | Image | Linewidth | Image | Linewidth |
| **10 µm** | 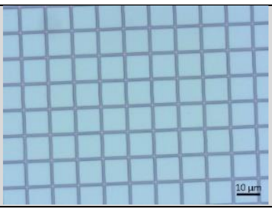 | 0.93 ± 0.55 | 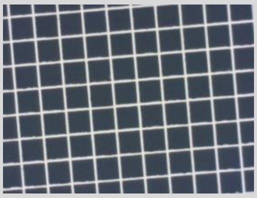 | 1.18 ± 0.11 |
| **22 µm** | 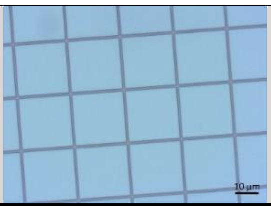 | 0.96 ± 0.05 | 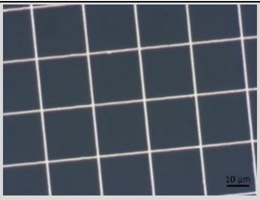 | 1.11 ± 0.06 |

#### S5 – PMG Sheet resistance vs. Gold Thickness – COMSOL Simulations

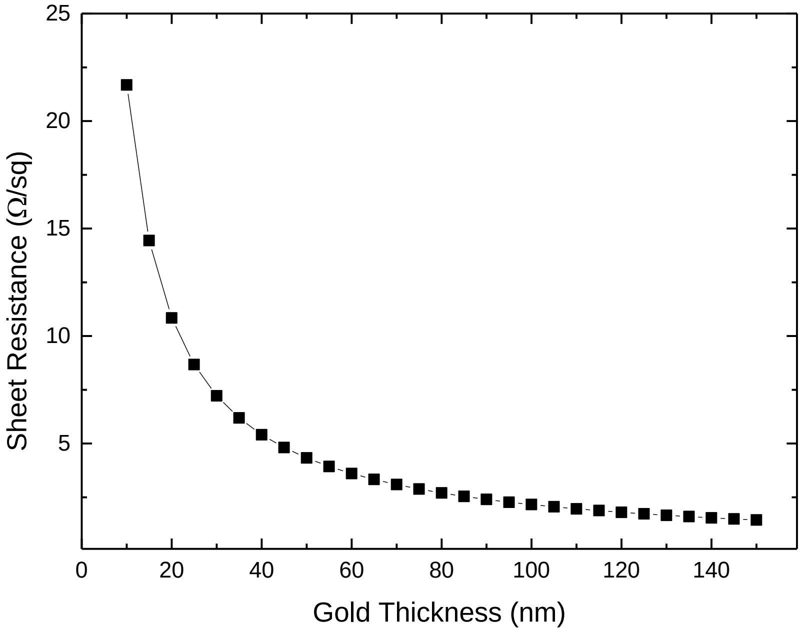


**Figure S5.** – PMG sheet resistance as a function of gold thickness, considering a spacing between grid lines of 10 μm and 1 μm linewidth. The present results were attained through COMSOL Multiphysics simulations.

### S6 – PMG Linewidth after Oxygen Plasma Pre-treatment

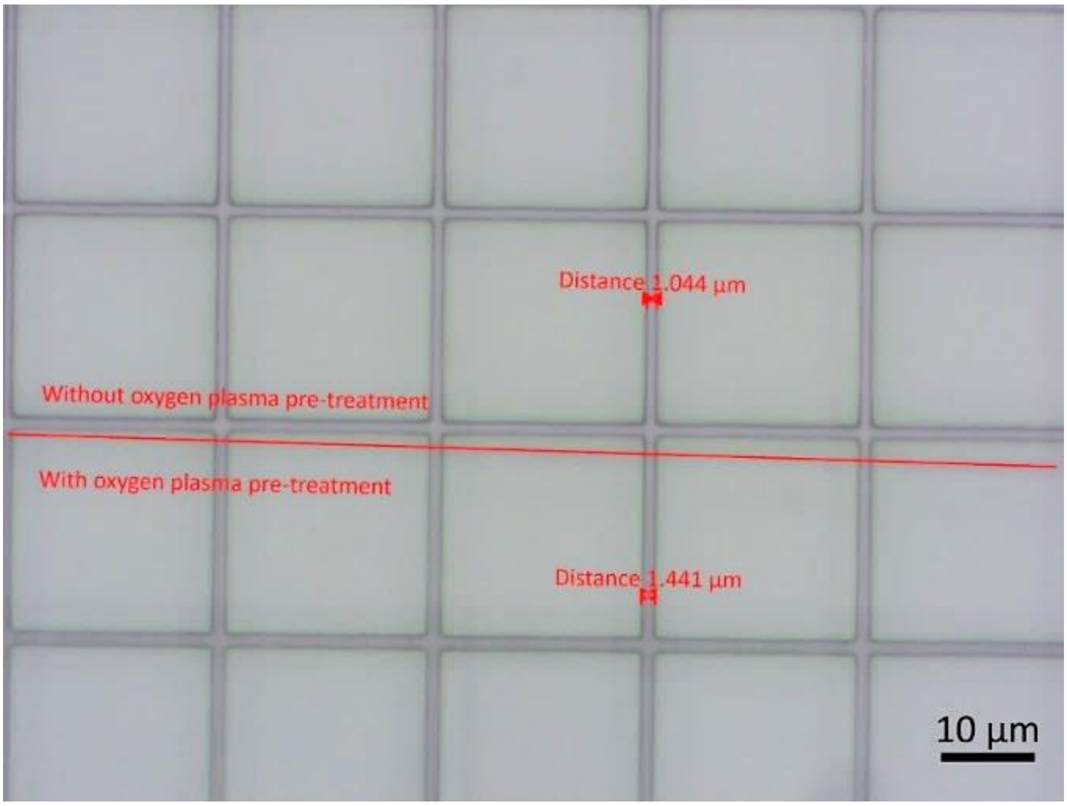


**Figure S6.** – Effect of oxygen plasma in the linewidth (1 µm) of patterned photoresist. The top half did not suffer any treatment. The bottom half was submitted to 5 minutes of oxygen plasma. The image was obtained through optical microscopy.

**S7 – Cytotoxic Effect Study Protocol (Materials and Methods)**

To evaluate if the electrodes substrate material could present toxicity once implanted in the body, cytotoxicity was evaluated in vitro according to the International Standard (ISO 10993–5) using the extract method.

Manipulations of Vero cells (African green Monkey kidney epithelial cells) and culture medium (DMEM Low Glucose w/ L-Glutamine w/ Sodium Pyruvate from Biowest, supplemented with 10% FBS (Fetal Bovine Serum) and 1% penicillin–streptomycin, from Invitrogen) were performed inside a biological safety cabinet (ESCO Lab- culture II). To produce the extracts, samples from the untreated and from the oxygen plasma treated material were sterilized with 70% v/v ethanol and, once fully dried, immersed the culture medium in the proportion of 6 mg of sample to 1 mL of medium and left for 48 h at 37 °C inside a 5% $CO_2$ humidified atmosphere incubator (Sanyo MCO19-AIC-UV). Cells from the Vero cell line were seeded at a density of 18k cells per $cm^2$ in a 96 well microplate (Sarstedt) and incubated in the $CO_2$ incubator for 24 h.

After that period, the medium was changed to proceed with the cytotoxicity assay. Four test conditions were considered: culture medium incubated for 48 h inside an incubator at 37 °C (negative control), conditioned medium by the untreated material, conditioned medium by oxygen plasma treated material, culture medium with 20% DMSO (Dimethyl sulfoxide) (positive control).

For each test condition four replicas were prepared using 100 μL of the respective medium. After 48 h of incubating the cells in these conditions, a viability test was performed. For this test, the media was replaced by a (1:1) v/v dilution in culture medium of a 0.04 g/mL resazurin solution in PBS (Phosphate buffered saline) (also used in control medium wells) and incubated for 3 h in the $CO_2$ incubator followed by absorbance readings at 600 nm and 570 nm (Biotek ELX 8000UV). The average values of 570 nm were subtracted to the average values of 600 nm, obtaining the correct average absorbance readings that are proportional to cell viability. Results are presented as cell viability relative to the negative control ± combined standard uncertainty calculated by propagation of the standard experimental deviations.
